# Individual-level metabolic connectivity captures cortical morphology and their coupling strengthens with age

**DOI:** 10.64898/2026.03.03.709267

**Authors:** Massimiliano Facca, Claudia Tarricone, Anna Ridolfo, Maurizio Corbetta, Andrei G. Vlassenko, Manu S. Goyal, Alessandra Bertoldo

**Affiliations:** Padova Neuroscience Center (PNC), University of Padova (Unipd), Padova, Italy; Department of Information Engineering, University of Padova (Unipd), Padova, Italy; Department of Neuroscience, University of Padova (Unipd), Padova, Italy; Venetian Institute of Molecular Medicine (VIMM), Padova, Italy; Neuroimaging Laboratories Research Center at the Mallinckrodt Institute of Radiology, Washington University School of Medicine, St Louis, MO, USA

**Keywords:** Brain Metabolism, FDG-PET, Metabolic Connectivity, Morphology, Ageing

## Abstract

**Purpose:** Cerebral glucose metabolism and cortical morphology are known to undergo significant changes across the lifespan, yet their network-level coordination remains poorly understood. This study aimed to investigate whether individual-level metabolic connectivity (MC) reflects underlying inter-areal morphometric similarity, and to determine how this metabolic–morphometric coupling evolves across the adult lifespan.

**Methods:** Dynamic [^18^F]FDG-PET and structural MRI data were acquired from 67 healthy adults (age range: 38–86 years). Individual MC networks were estimated based on the similarity between regional time–activity curves. Corresponding structural similarity networks were generated using the morphometric inverse divergence (MIND) framework, which integrates multiple vertex-wise features of cortical morphology. The correspondence between metabolic and structural networks was quantified at both global and local scales using Spearman correlations. General linear models were employed to assess age-related effects on MC–MIND similarity.

**Results:** MC demonstrated a robust positive association with cortical morphometric similarity (ρ = 0.32, p < 0.0001), an association that persisted after distance correction and was replicated at the individual level. Regional coupling followed a topographic gradient, peaking in heteromodal association cortices and reaching its minimum in paralimbic areas. Crucially, morphology–metabolism alignment systematically strengthened with age at the global level (β = 0.59, p < 0.001). Local age-related increases were spatially heterogeneous, predominantly affecting visual, dorsal parietal, and premotor cortices alongside adjacent multimodal regions.

**Conclusion:** Individual-level MC captures the morphometric organisation of the brain. The age-related increase in morphology–metabolism coupling indicates that metabolic coordination becomes progressively more aligned with cortical architecture, consistent with reduced neuroenergetic flexibility in the ageing brain.

## Introduction

The human brain functions as a complex network of interconnected regions that continuously coordinate their activity to support perception, cognition, and behaviour. Elucidating how these distributed elements communicate and integrate information has long been a central endeavour in neuroscience [1]. Network-based analyses have revealed that the brain’s architecture is governed by consistent organisational principles—modularity, hierarchy, and small-world topology—whose interplay strikes a balance between functional specialisation and efficient global integration [2, 3]. Functional and structural connectivity mapping with Magnetic Resonance Imaging (MRI) has been pivotal in uncovering these principles, delineating how neural systems interact dynamically and how such interactions are related to anatomical pathways [4–6].

Whereas functional MRI captures transient fluctuations in neuronal activity and diffusion MRI delineates the architecture of white-matter tracts, metabolic imaging offers a complementary window onto the brain’s energetic landscape [7]. Because neuronal signalling constitutes the predominant driver of cerebral glucose consumption [8], coupling in glucose utilisation reflects coordinated energetic demands underlying large-scale brain dynamics [9, 10]. This coordination can be quantified through metabolic connectivity, an energetic analogue of functional connectivity that grounds network organisation in the bioenergetic processes sustaining neural activity [10, 11]. Often, the metabolic signature of brain networks is inferred from static [^18^F]FDG-PET acquisitions, in which inter-regional correlations of semiquantitative uptake measures—typically standardised uptake values (SUVs)—are computed across subjects [12–15]. This population-level approach has yielded valuable insights into metabolic network disruptions across neurological and psychiatric disorders [16, 17], yet it remains inherently limited to the group level, as inter-regional metabolic associations are derived from across-subject covariance matrices rather than intra-individual dynamics, and cannot naturally capture interindividual variability in metabolic interactions, thereby restricting its applicability for precision phenotyping. Recently, new perspectives have emerged within the molecular imaging field, with increasingly sophisticated frameworks enabling metabolic connectivity to be delineated at the single-subject level [18, 19]. The development of these approaches has been fostered not only by dynamic [^18^F]FDG-PET acquisitions made via conventional bolus protocols, but also through the implementation of constant infusion and hybrid bolus–infusion designs [20–23].

Metabolic connectivity derived from dynamic [^18^F]FDG-PET captures similarity in glucose utilisation across distributed brain regions, reflecting coordinated metabolic demands across neural populations [24]. Metabolic connectivity has shown non-redundancy with functional connectivity, along with potentially heightened sensitivity to physiological alterations [25, 26], as well as a closer association with cognitive processes [26, 27]. However, the interpretative value of this measure ultimately hinges on its biological grounding—that is, on how patterns of metabolic coupling relate to other fundamental dimensions of brain organisation, including structural connectivity, cortical morphology, and neurochemical architecture. Establishing these relationships is crucial for determining what metabolic connectivity reveals about the underlying neural substrate and for assessing its sensitivity to physiological processes as well as to pathological alterations observed in neurodegenerative and neoplastic diseases [25, 28, 29].

To address this question, we focus on the relationship between metabolic connectivity and cortical morphology—a key structural dimension that captures the brain’s cytoarchitectonic organisation. Recent advances in structural neuroimaging have enabled increasingly precise quantification of cortical morphology, moving beyond univariate descriptors such as cortical thickness or surface area towards network-based representations, thereby yielding structural similarity networks [30]. Among these, the morphometric inverse divergence (MIND) [31] framework provides a principled approach to quantify structural similarity between cortical regions by comparing multivariate distributions of morphological features. Networks constructed using MIND recapitulate canonical principles of cortical organisation—linking cytoarchitectonically similar, homologous, and axonally connected regions—and show strong correspondence with transcriptional similarity patterns across the human cortex [31]. On this basis, we hypothesised that similarity in cortical morphology would be associated with similarity in kinetic profiles captured by metabolic connectivity. Because morphometric similarity reflects shared cytoarchitectonic organisation, regions with comparable profiles may operate under similar energetic constraints [8], which is consistent with the principle of structural homophily whereby regions with similar structural properties tend to exhibit coordinated functional dynamics [32].

Here, we integrate dynamic [^18^F]FDG-PET and structural MRI data acquired from the same individuals to investigate how cortical morphology constrains large-scale patterns of metabolic connectivity across the human lifespan. By combining structural and metabolic information from the same brains, we demonstrate a global alignment between metabolic connectivity and morphological similarity, which is evident at both the group and individual levels and follows a spatial gradient rather than a uniform cortical distribution. Notably, this correspondence systematically increases with age, suggesting that metabolic dynamics become progressively more constrained by the brain’s structural architecture through adulthood, both globally and in a region-specific fashion. Collectively, these findings indicate that shared microarchitectural organisation provides a structural and energetic scaffold for coherent metabolic dynamics.

## 2 Material and methods

### 2.1 Research participants

The study cohort comprised 67 healthy individuals (35 females; mean age = 63.8 ± 14.6 years, range = 37.9–86.3 years). All imaging procedures were approved by the Human Research Protection Office and the Radioactive Drug Research Committee at Washington University in St. Louis.

### 2.2 PET and MRI data acquisition

PET data were acquired on a Siemens Biograph Vision scanner (Siemens Healthineers, Erlangen, Germany). An intravenous catheter was placed for tracer administration, and [^18^F]-fluorodeoxyglucose (FDG) was administered as a single bolus injection. The mean injected dose was 197.65 ± 14.21 MBq. The head position was stabilised to minimise motion throughout the scan. Dynamic data were reconstructed from list-mode acquisitions via a three-dimensional (3D) ordered-subset expectation maximisation algorithm with time-of-flight information (OSEM+TOF; 8 iterations, 5 subsets), resulting in 220 × 220 × 159 volumes with an isotropic voxel size of approximately 1.65 mm. The dynamic acquisition consisted of 52 frames with increasing durations: 10 × 6 s, 8 × 15 s, 9 × 60 s, and 12 × 240 s. Standard corrections for radioactive decay, attenuation, scatter, and random coincidences were applied. No post-reconstruction spatial smoothing was performed.

MRI data were acquired on a 3T Siemens Magnetom Prisma scanner. Structural images were obtained using a 3D sagittal T1-weighted MPRAGE sequence with four echoes (TE = 1.81, 3.60, 5.39, and 7.18 ms), TR = 2500 ms, TI = 1000 ms, and flip angle = 8°. Images were acquired at 0.8 mm isotropic resolution.

### 2.3 Preprocessing of PET and MRI data

#### 2.3.1 Preprocessing of T1w data

For each subject, a T1w image was generated by averaging the first two echoes of the multi-echo MPRAGE acquisition. Brain extraction, segmentation, and surface reconstruction were performed using FreeSurfer (v7.4.1) [33]. Individual surfaces were subsequently registered to the fsLR standard surface space using the Connectome Workbench [34]. One participant was excluded from the final sample due to technical failures during the FreeSurfer *recon-all* procedure, resulting in a final cohort of 66 individuals.

#### 2.3.2 Preprocessing of PET data

Dynamic [^18^F]FDG-PET data were preprocessed using an in-house pipeline integrating FreeSurfer, FSL, Connectome Workbench, and SynthStrip [35-37]. Motion correction was implemented in two stages. In the first stage, frames acquired after 2 minutes post-injection were smoothed (6 mm FWHM Gaussian kernel) and aligned to the earliest frame exceeding this cutoff using FSL’s *flirt*. The motion-corrected frames were then summed to generate a preliminary reference image. In the second stage, all frames were re-aligned to this refined reference. The resulting motion-corrected images were summed again to produce a second template, which was skull-stripped using SynthStrip and rigidly registered to the individual T1w image. Motion-correction and registration transformations were concatenated and subsequently applied to the original, unsmoothed data to achieve simultaneous motion correction and resampling into T1w space. Following this step, PET data were projected onto each participant’s cortical surface and resampled onto the fsLR (32k) standard surface using the Connectome Workbench. Vertex-wise time–activity curves (TACs) were then averaged according to the Glasser atlas [38], yielding 360 regional TACs. No spatial smoothing was applied to the PET data. Additionally, as a supplementary control, dynamic PET data were corrected for partial volume effects (PVE) using the iterative Yang method [39] implemented in PETPVC [40]. The tissue maps for PVC were based on FreeSurfer’s GTM segmentation, which includes both cerebral and extra-cerebral structures. The resulting corrected data were then projected onto the cortical surface, parcellated as described above, and used to evaluate the robustness of our findings to PVC.

### 2.4 Construction of Morphometric Similarity Networks

Structural similarity networks were derived using the morphometric inverse divergence (MIND) framework [31], a general approach for estimating regional similarity based on the multivariate Kullback–Leibler (KL) divergence between morphometric feature distributions. MIND quantifies how closely two regions resemble each other by comparing their full multivariate morphometric profiles using a k-nearest-neighbour estimator. The resulting divergence values are inverted and linearly rescaled, such that higher similarity corresponds to values closer to 1, whereas larger divergences yield lower scores. MIND matrices were constructed according to the Glasser parcellation, yielding symmetric 360 × 360 matrices. As in the original implementation, MIND was applied to vertex-wise morphometric measures—cortical thickness, mean curvature, sulcal depth, surface area, and volume—extracted using FreeSurfer. Full methodological details are provided in the original work of Sebenius et al. (2023) [31].

### 2.5 Metabolic connectivity mapping

Individual-level metabolic connectivity matrices were derived using a distance-based similarity framework [19]. For each pair of regions, the coupling strength was defined as one minus the Euclidean distance between their [^18^F]FDG-PET TACs, after scaling distances by the maximum value across all pairs to constrain results to the [0, 1] range. In this representation, higher values reflect greater pharmacokinetic similarity in regional [^18^F]FDG uptake.

### 2.6 Statistical analyses

### 2.6.1 Group-level correspondence

Group-level MIND and MC matrices were obtained by averaging individual matrices across participants. The global correspondence between modalities was quantified by computing the Spearman correlation between the vectorised triangle of the two group-level matrices. To verify that this correspondence was not driven by inter-regional distance, we regressed out spatial proximity effects. For both MIND and MC, edge weights were modelled as a function of the Euclidean distance (d) between ROI centroids using a nonlinear exponential decay model of the form *w*(*d*)= *ae*^−*bd*^ + *c*. Model parameters were estimated using nonlinear least-squares optimisation and the residuals from these fits were used to recompute the global Spearman correlation. We then examined whether regions that were more strongly connected in one modality also showed higher connectivity in the other. For each ROI, weighted degree (nodal strength) was computed as the sum of its edge weights. Correspondence between modalities was assessed by computing the Pearson correlation between nodal strength maps. Statistical significance was evaluated while accounting for spatial autocorrelation using a non-parametric spin test [41] (N spins = 10,000).

Local correspondence between modalities was quantified by computing, for each ROI, the Spearman correlation between its MC profile and its MIND profile (i.e., between matched rows of the two matrices). These coefficients (ρ) indexed regional MC–MIND coupling. The spatial distribution of ρ values was then compared across Mesulam’s laminar classes [42]. Deviations from chance were assessed using spin tests to control for spatial autocorrelation (N spins = 10,000).

#### 2.6.2 Individual-level coupling

To evaluate whether group-level correspondence was preserved at the individual level, analyses were repeated within subjects. To test whether the MC–MIND relationship reflected subject-specific coupling, a permutation-based null model was constructed by randomly reassigning MIND matrices across participants, such that each subject’s MC matrix was paired with a MIND matrix from a different individual (1,000 permutations). Global MC–MIND similarity was recomputed for each mismatched pair and averaged across participants, yielding a null distribution of mean correlations. The empirical p-value was defined as the proportion of permuted means exceeding the observed mean correlation across subjects. The same procedure was repeated after regressing out inter-regional distance effects from both MC and MIND matrices, following the group-level analyses, to verify that subject-specific coupling was not driven by spatial proximity.

#### 2.6.3 Effects of age on global and local coupling

To examine age-related effects on MC–MIND similarity, both global and regional indices were entered into general linear models (GLMs) with age, sex, and total intracranial volume (TIV) as predictors. Continuous predictors and response variables were z-scored prior to the analysis. A bootstrap resampling procedure was performed to evaluate the stability of the associations, with 10,000 resamples used to derive 95% confidence intervals for the beta coefficients. For local (i.e., regional) coupling analyses, multiple comparisons were controlled using false discovery rate (FDR) correction across regions. Effect sizes are reported as partial correlations (i.e., adjusted for age and sex), and only effects surviving FDR correction were considered statistically significant.

## 3 Results

### 3.1 Global correspondence between metabolic connectivity and morphometric similarity

We first assessed how strongly MC aligns with morphometric similarity at the global level (Fig. 1a,b). The two group-level matrices showed a clear positive correspondence (ρ = 0.32, p < 0.0001; Fig. 1c), which remained significant after controlling for inter-regional distance (ρ_dist-corr_ = 0.31, p < 0.0001), indicating that the relationship was not driven by spatial proximity. A similar pattern was observed at the individual level, with consistent positive MC–MIND coupling across participants (mean ρ = 0.28 ± 0.03; range = 0.23–0.34; all p < 0.0001; Fig. 1d). Within-subject MC–MIND coupling remained significant after regressing out inter-regional distance (mean ρ_dist-corr_ = 0.27 ± 0.03; range = 0.22–0.34; all p < 0.0001; Fig. 1d). We next examined correspondence in regional connectivity strength. MIND and MC nodal strength values were positively correlated at the group level (r = 0.42, p_spin_ < 0.01; Fig. 1e), indicating that regions exhibiting higher morphometric similarity to the rest of the cortex also tend to show stronger metabolic integration. This strength-based relationship was replicated at the individual level (mean r = 0.40 ± 0.03; range = 0.36–0.54; all p_spin_ < 0.05). Finally, we tested whether MC–MIND coupling exhibited subject specificity beyond chance. Empirical within-subject correlations consistently exceeded those obtained from the null distribution (p < 0.001), indicating that the observed subject-specific correspondence was greater than expected under random pairing.

**Fig. 1.**
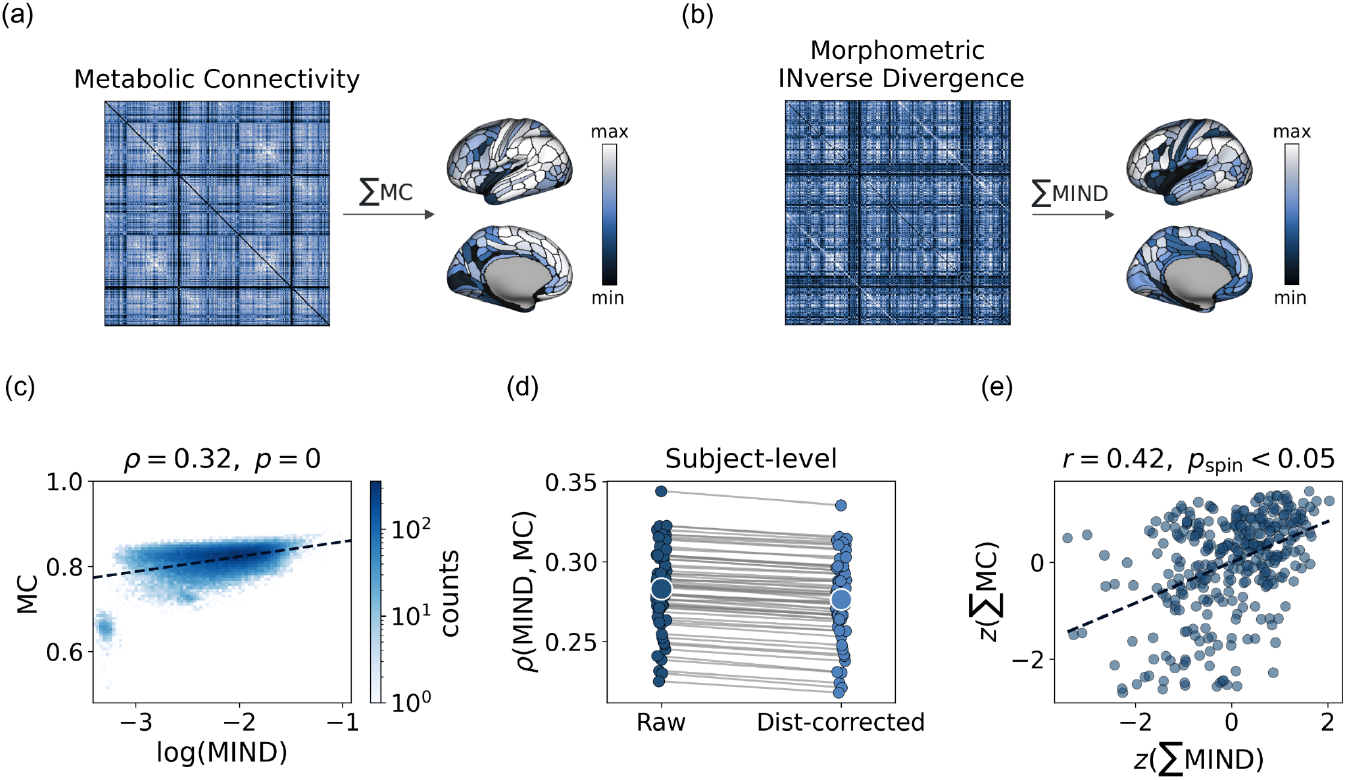
Overview of the data and global relationships between metabolic connectivity and morphometric similarity. (a) Group-averaged metabolic connectivity (MC) matrix together with the corresponding regional weighted degree map. (b) Group-averaged morphometric inverse divergence (MIND) matrix and associated regional weighted degree. (c) Edge-wise association between MC and MIND, with a quantile (median) regression line overlaid for visualisation. MIND values were log-transformed for display purposes. The colorbar indicates the density of points. (d) Edge-wise association between MC and MIND at the single-subject level, before and after correcting both MC and MIND for distance-dependent effects via exponential fitting. Each dot represents the MC–MIND similarity value for a single subject. Similarity measures decreased only slightly, indicating that the coupling between MC and MIND is robust to distance effects. (e) Pearson correlation between the regional weighted-degree vectors of MIND and MC; significance was assessed using a spin test to account for spatial autocorrelation

### 3.2 The relationship between metabolic connectivity and morphometric similarity increases with age

We then evaluated whether interindividual differences in MC–MIND coupling were associated with age. Global coupling increased significantly with age (β = 0.59, p < 10^−6^) after controlling for sex and TIV (Fig. 2). A bootstrap resampling analysis further supported the stability of this association, yielding a 95% confidence interval of [0.39, 0.79]. No significant relationship was observed for sex (p = 0.17). Residual diagnostics indicated no evidence of autocorrelation or major violations of model assumptions (Supplementary Fig. S1). Together, these findings indicate that the correspondence between MC and MIND strengthens with age, suggesting that metabolic dynamics become progressively more constrained by the brain’s anatomical architecture.

**Fig. 2.**
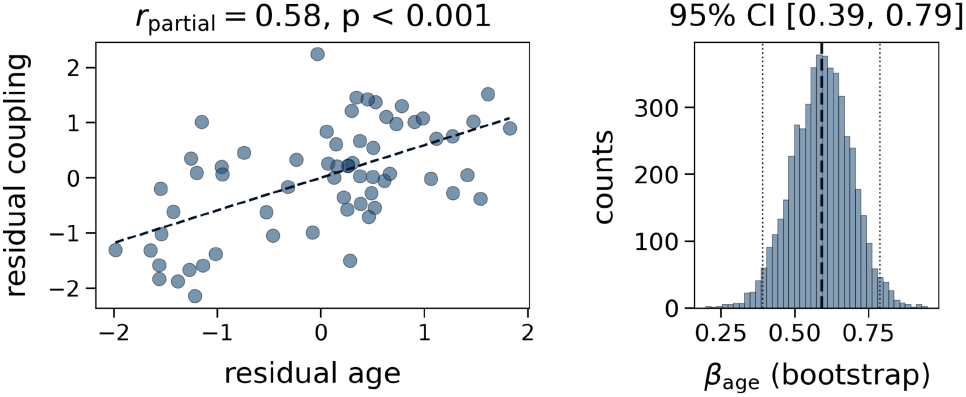
Global coupling between metabolic connectivity and morphometric similarity increases with age. Scatterplot showing the relationship between age and the global coupling between MC and MIND, after both variables were residualised for sex and total intracranial volume (TIV). The corresponding partial correlation is reported. The right panel displays the confidence interval (CI) for β_age_

### 3.3 Spatially structured correspondence between MC and MIND

We then characterised the regional distribution of MC–MIND coupling across the cortex. Local correspondence exhibited a structured topographic pattern (Fig. 3a). Coupling values were highest in dorsal frontoparietal regions, including intraparietal cortex and lateral prefrontal areas, which showed the greatest similarity between MC and MIND. In contrast, markedly lower values were observed in medial paralimbic territories and archicortical structures, including the anterior cingulate cortex, ventromedial prefrontal cortex, and hippocampal–presubicular areas, where correspondence between metabolic and structural similarity was comparatively weak (Fig. 3a). A distinct spatial pattern characterised the median absolute deviation (MAD) of local coupling across subjects: variability peaked in dorsal parietal and intraparietal regions, whereas orbitofrontal and anterior temporal areas exhibited comparatively low dispersion (Fig. 3b). Quantitatively, MC–MIND coupling aligned with established large-scale organisational axes. Across Mesulam’s laminar classes, coupling was strongest in the heteromodal association cortex (p_spin_ < 0.05; Fig. 3c) and reduced in paralimbic regions (p_spin_ < 0.05; Fig. 3c). Together, these findings indicate that alignment between metabolic and morphometric organisation follows a structured spatial pattern rather than being uniformly distributed across the cortex.

**Fig. 3.**
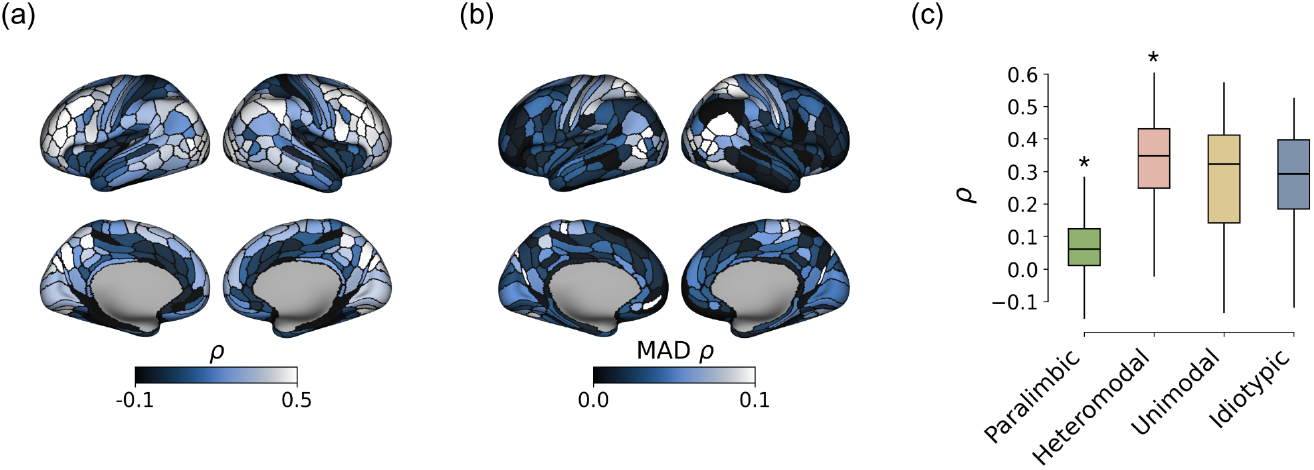
Local coupling between MC and MIND. (a) Regional variability of local MC–MIND coupling, computed as the Spearman correlation between corresponding node-wise connectivity profiles. (b) Median absolute deviation (MAD) of local coupling across subjects. (c) Distribution of local MC–MIND coupling across Mesulam’s laminar classes, with statistical significance assessed using spin tests. Asterisks (*) indicate statistically significant deviations—either higher or lower than expected by chance

### 3.4 Age-related variability in local metabolic - morphological relationships

We then tested whether age modulated the regional correspondence between MC and MIND. Age exerted a widespread influence across the cortex, with 102 out of 360 regions surviving FDR correction (Fig. 4). Nearly all the significant regions (98%) showed positive associations, with partial correlations ranging from 0.30 to 0.63. Age-related increases in local coupling displayed a clear spatial structure, with effects concentrated in early and higher-order visual areas, the dorsal parietal cortex, premotor and motor territories, and portions of the temporo-parieto-occipital junction. Additional significant associations were observed in lateral temporal regions, parahippocampal areas, and selected insular territories, indicating that the age-related strengthening of MC–MIND correspondence extends beyond the canonical dorsal and ventral visual pathways to include adjacent multimodal cortices. These spatial patterns suggest that these areas are particularly sensitive to age-associated shifts towards morphology-constrained metabolic organisation. No region showed a significant effect of sex after FDR correction (all p > 0.05).

**Fig. 4.**
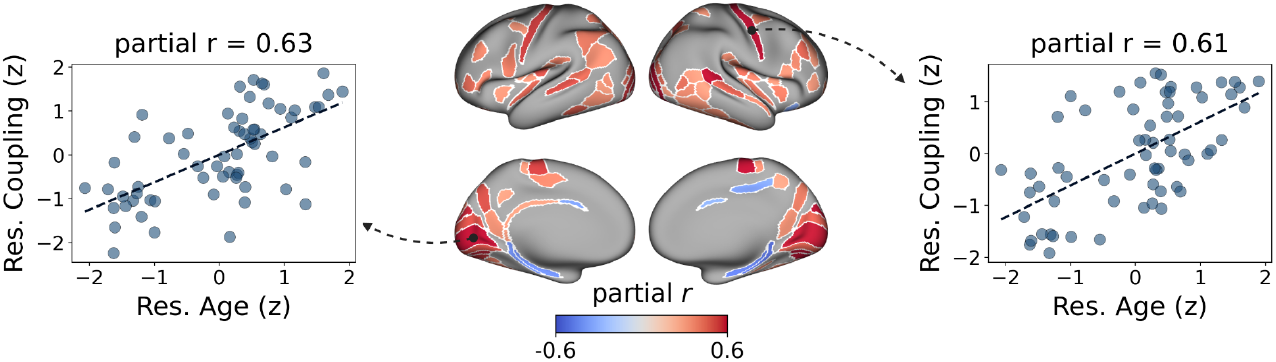
Age effects on local coupling between MC and MIND. Regional distribution of ROIs showing a significant relationship (FDR-corrected) between local MC–MIND coupling and age, after controlling for sex and TIV. The effect of age on coupling was positive in 98% of the ROIs. The leftmost and rightmost panels show two example correlations

### 3.5 Robustness of findings to partial volume effect correction

To ensure that the observed relationship between MC and MIND was not driven by partial volume effects, we replicated our primary analyses using PVE-corrected PET data. The global correspondence between metabolic and morphometric organisation remained highly significant and was slightly stronger than that observed in the primary analysis (ρ = 0.35, p < 0.0001). This consistency was also evident at the individual level (mean ρ = 0.29 ± 0.03; range = 0.21–0.33, all p < 0.0001). Notably, the age-related increase in global coupling was preserved after PVE correction, resulting in an even larger effect size than that in the primary model (*β* = 0.65, p < 10^−8^; see Supplementary Fig. S2). At the regional level, the number of significant regions, the direction of the associations, and the overall effect sizes for age-related changes in local coupling remained largely consistent with the main findings (Supplementary Fig. S3).

## 4 Discussion

In this study, we combined dynamic [^18^F]FDG-PET with structural MRI to examine how cortical morphology shapes large-scale patterns of metabolic connectivity across the adult lifespan. We assessed this relationship at multiple levels—global, individual, and regional—and found a robust positive association between the two networks. The correspondence persisted after accounting for spatial distance, was consistently detectable in individual participants, displayed regional heterogeneity across canonical cortical systems, and systematically strengthened with advancing age.

The observed correspondence between MC and cortical morphology indicates a fundamental coupling between structural architecture and metabolic coordination in the human brain. Regions sharing similar morphometric profiles—likely reflecting convergent cytoarchitectonic organisation—also display coherent temporal patterns of glucose utilisation. This relationship likely stems from shared cellular determinants of energy demand, including neuronal and synaptic density, myelination, and glial composition [8]. Within this framework, the metabolic coupling captured by MC can be viewed as an energetic expression of structural homophily: regions with analogous cytoarchitectonic profiles may operate under comparable biophysical and vascular constraints, supporting the optimisation of energy use across the cortex while maintaining coordinated signalling among structurally similar territories. Importantly, this alignment may also reflect shared transcriptional programmes that govern cortical differentiation and maturation [30, 32, 43]. Genes involved in neuronal growth, synaptic organisation, and myelination follow coordinated spatial and temporal trajectories during development, potentially giving rise to enduring similarities in both microarchitecture and metabolic function [32, 43]. This view aligns with reports that regions with similar transcriptional profiles tend to exhibit comparable structural and functional connectivity, as well as related cytoarchitectonic features [44–46]. However, the fact that this alignment is moderate suggests a degree of structural–metabolic decoupling. While morphology provides a stable biological scaffold, it does not strictly determine brain metabolic coordination. This divergence likely reflects a degree of neuroenergetic flexibility, which may be grounded in the different timescales governing these two modalities. Morphometric similarity is a relatively stable architectural feature that evolves over months or years, reflecting slow processes of structural remodelling. In contrast, metabolic connectivity, as captured by regional glucose kinetics, reflects energetic demands that can be reconfigured over much shorter periods (i.e., minutes) to support transient physiological states. This temporal mismatch implies that the metabolic network maintains a degree of autonomy from the cortical architecture, enabling a dynamic repertoire of energetic states that are not bound by the slower pace of structural change.

Another major finding of this work is the progressive strengthening of the coupling between MC and structural similarity with age. This pattern, observed across global and regional scales, suggests that the coordination of glucose metabolism becomes increasingly governed by cortical morphology over the adult lifespan. Age-related remodelling of microarchitecture—including dendritic simplification, synaptic pruning, and myelin degeneration—occurs alongside reductions in mitochondrial efficiency and glycolytic capacity [47–49]. Rather than indicating a uniform metabolic decline, our findings point to a reorganisation of neuroenergetic coupling. This trend likely reflects a gradual loss of metabolic flexibility, consistent with fMRI evidence showing age-related increases in structure–function coupling and reduced network reconfiguration capacity [50]. This interpretation is further supported by recent fPET evidence indicating that ageing is associated with reduced temporal variability (standard deviation) in regional glucose uptake dynamics [51]. While Deery and colleagues [51] focused on the inherent fluctuations of the metabolic signal, our results complement their findings by showing that this loss of dynamic range is accompanied by a tighter alignment with the underlying anatomy. Together, these observations suggest that the ageing brain undergoes a shift towards a more “static” metabolic organisation: as the temporal variability of glucose consumption decreases, its spatial pattern becomes increasingly anchored to the physical substrate. Importantly, we extended our analyses to the local level and identified widespread age-related increases in morphology–metabolism alignment across sensory, motor, and transmodal cortices. This suggests that the transition towards a more rigid, structure-bound metabolic organisation is a global hallmark of the ageing brain, characterised by a reduced capacity for energetic reconfiguration across multiple functional systems. Finally, the stability of our findings following PVE correction is a critical methodological point. Given that ageing is inherently associated with cortical thinning—which can exacerbate partial volume effects—ensuring that the observed MC–MIND coupling was not a byproduct of anatomical atrophy was essential. The fact that the age-related effects were preserved, and even strengthened, after correction validates the increasing metabolic–structural convergence as a physiological shift characterising the ageing brain.

From a clinical standpoint, quantifying the correspondence between cortical morphology and glucose metabolism may provide a sensitive index of neuroenergetic efficiency and structural integrity across the lifespan. Because [^18^F]FDG-PET is widely used in clinical neurology, the association between metabolic connectivity and morphological similarity could offer a complementary descriptor of brain organisation beyond regional uptake measures. In neurodegenerative diseases such as Alzheimer’s disease and Parkinson’s disease, alterations in metabolic connectivity are frequently observed [52–55]. Evaluating structure–metabolism coupling at the network level may help determine whether these metabolic abnormalities reflect disrupted energetic coordination or emerge secondarily from deteriorating structural relationships among cortical areas, for example, owing to regional atrophy. Similarly, in oncological conditions, tumour infiltration may locally alter cortical tissue properties in ways that could disrupt the spatial coherence of glucose metabolism [29]. Characterising such alterations in terms of structure–metabolism coupling may enhance the interpretability of metabolic imaging by clarifying how tissue changes propagate to large-scale metabolic interactions. Finally, longitudinal assessment of morphology–metabolism coupling could provide a complementary marker of disease progression, particularly in conditions where metabolic dysregulation and structural decline evolve at different rates.

### Limitations

Several methodological considerations should be noted. First, dynamic [^18^F]FDG-PET with a bolus protocol characterises inter-regional coupling based on similarities in regional kinetic profiles, rather than rapid metabolic fluctuations, which are typically assessed using fPET protocols [56]. Although this approach yields coarser temporal resolution, it offers a higher signal-to-noise ratio and well-established quantification procedures, making it suitable for estimating metabolic connectivity in standard clinical acquisitions. Second, although morphometric similarity provides a biologically grounded representation of cortical structure, it does not incorporate subcortical structures or white matter pathways, which may also contribute to large-scale metabolic coordination. A key advantage of this approach over tractography-based connectivity lies in its ability to capture structural correspondence across all cortical regions—including those not directly linked by white matter tracts [30]. This makes it a suitable framework for studying metabolic connectivity, which reflects coordinated dynamics that may arise between regions that are not anatomically adjacent but coupled through multistep pathways. Finally, the cohort consisted of participants older than 35 years. While this limits generalisability to younger populations, it focuses on an age range in which age-related and neurodegenerative processes become increasingly prominent, making it particularly relevant for investigating structure–metabolism coupling in the ageing brain.

### Conclusions

Taken together, these findings demonstrate that cortical morphology and metabolic connectivity are interrelated, and that their coupling systematically strengthens with age. Given the widespread clinical use of [^18^F]FDG-PET, quantifying morphology–metabolism correspondence may provide a translational marker of neuroenergetic integrity. Such an approach could improve the interpretation of metabolic alterations in ageing and disease, help distinguish primary metabolic dysregulation from structural deterioration, and ultimately contribute to more refined, multimodal phenotyping in clinical neuroimaging.

## Acknowledgements

We appreciate the members of the Vlassenko-Goyal Laboratory (VG Lab), Center for Clinical Imaging Research (CCIR), the Knight Alzheimer Disease Research Center, and the Cyclotron Facility at Washington University for making this research possible. We are particularly grateful to our research participants for their altruism. Some of the MRI sequences used in the CCIR were obtained from the Massachusetts General Hospital.

## Declarations

### Author Contributions

**M.F**.: conceptualisation, methodology, software, formal analysis and investigation, writing - original draft. **C.T**.: writing - review and editing. **A.R**.: writing - review and editing. **M.C**.: writing - review and editing. **A.G.V**.: data collection, writing - review and editing. **M.S.G**.: data collection, writing - review and editing. **A.B**.: supervision, methodology, writing - review and editing.

### Funding

MF, CT, AR, MC and AB received support for the research, authorship, and publication of this article by the European Union under the project *EBRAINS 2*.*0: A Research Infrastructure to Advance Neuroscience and Brain Health* (HORIZON-INFRA-2022-SERV-B-01, grant no. 101147319). AGV and MSG received support for this research from the National Institute on Aging of the National Institutes of Health, U.S.A. (NIA/NIH R01AG073210, R01AG057536, and R01053503).

### Ethics approval

This study was performed in line with the principles of the Declaration of Helsinki. Approval was granted by the Human Research Protection Office and the Radioactive Drug Research Committee at Washington University in St. Louis.

### Consent to participate

Informed consent was obtained from all individual participants included in the study.

### Competing interests

The authors declare no competing interests.

### Data availability

The imaging data used in this study are available upon reasonable request by a qualified researcher following a data use agreement. Requests for imaging data should be directed to the VG Lab.

## Notes

### Competing Interest Statement

The authors have declared no competing interest.

## References

1. Barabási DL, Bianconi G, Bullmore E, et al. Neuroscience Needs Network Science. J Neurosci. 2023;43(34):5989–5995. 10.1523/JNEUROSCI.1014-23.2023

2. Bassett DS, Sporns O. Network neuroscience. Nat Neurosci. 2017;20(3):353–364. 10.1038/nn.4502

3. Bullmore E, Sporns O. Complex brain networks: graph theoretical analysis of structural and functional systems. Nat Rev Neurosci. 2009;10(3):186–198. 10.1038/nrn2575

4. Fotiadis P, Parkes L, Davis KA, et al. Structure-function coupling in macroscale human brain networks. Nat Rev Neurosci. 2024;25(10):688–704. 10.1038/s41583-024-00846-6

5. Park HJ, Friston K. Structural and functional brain networks: from connections to cognition. Science. 2013;342(6158):1238411. 10.1126/science.1238411

6. Suárez LE, Markello RD, Betzel RF, Misic B. Linking Structure and Function in Macroscale Brain Networks. Trends Cogn Sci. 2020;24(4):302–315. 10.1016/j.tics.2020.01.008

7. Sala A, Lizarraga A, Caminiti SP, et al. Brain connectomics: time for a molecular imaging perspective? Trends Cogn Sci. 2023;27(4):353–366. 10.1016/j.tics.2022.11.015

8. Attwell D, Iadecola C. The neural basis of functional brain imaging signals. Trends Neurosci. 2002;25(12):621–625. 10.1016/s0166-2236(02)02264-6

9. Horwitz B, Duara R, Rapoport SI. Intercorrelations of glucose metabolic rates between brain regions. J Cereb Blood Flow Metab. 1984;4(4):484–499. 10.1038/jcbfm.1984.73

10. Watabe T, Hatazawa J. Evaluation of Functional Connectivity in the Brain Using PET: A Mini-Review. Front Neurosci. 2019;13:775. 10.3389/fnins.2019.00775

11. Reed MB, Cocchi L, Sander CY, et al. Connecting the dots: approaching a standardized nomenclature for molecular connectivity in PET. Eur J Nucl Med Mol Imaging. 2025. 10.1007/s00259-025-07357-1

12. Di X, Biswal BB, ADNI. Metabolic brain covariant networks as revealed by FDG-PET with reference to resting-state fMRI networks. Brain Connect. 2012;2(5):275–283. 10.1089/brain.2012.0086

13. Yakushev I, Chételat G, Fischer FU, et al. Metabolic and structural connectivity within the DMN relates to working memory. Neuroimage. 2013;79:184–190. 10.1016/j.neuroimage.2013.04.069

14. Yakushev I, Ripp I, Wang M, et al. Mapping covariance in brain FDG uptake to structural connectivity. Eur J Nucl Med Mol Imaging. 2022;49(4):1288–1297. 10.1007/s00259-021-05590-y

15. Savio A, Fünger S, Tahmasian M, et al. Resting-State Networks as Simultaneously Measured with fMRI and PET. J Nucl Med. 2017;58(8):1314–1317. 10.2967/jnumed.116.185835

16. Caminiti SP, Tettamanti M, Sala A, et al. Metabolic connectomics targeting brain pathology in dementia with Lewy bodies. J Cereb Blood Flow Metab. 2017;37(4):1311–1325. 10.1177/0271678X16654497

17. Huang S, Li J, Sun L, et al. Learning brain connectivity of Alzheimer’s disease by sparse inverse covariance estimation. Neuroimage. 2010;50(3):935–949. 10.1016/j.neuroimage.2009.12.120

18. Jamadar SD, Ward PGD, Liang EX, et al. Metabolic and Hemodynamic Resting-State Connectivity of the Human Brain. Cereb Cortex. 2021;31(6):2855–2867. 10.1093/cercor/bhaa393

19. Volpi T, Vallini G, Silvestri E, et al. A new framework for metabolic connectivity mapping using bolus [18F]FDG PET and kinetic modeling. J Cereb Blood Flow Metab. 2023;43(11):1905–1918. 10.1177/0271678X231184365

20. Hahn A, Gryglewski G, Nics L, et al. Quantification of Task-Specific Glucose Metabolism with Constant Infusion of 18F-FDG. J Nucl Med. 2016;57(12):1933–1940. 10.2967/jnumed.116.176156

21. Li S, Jamadar SD, Ward PGD, et al. Analysis of continuous infusion functional PET (fPET) in the human brain. Neuroimage. 2020;213:116720. 10.1016/j.neuroimage.2020.116720

22. Villien M, Wey HY, Mandeville JB, et al. Dynamic functional imaging of brain glucose utilization using fPET-FDG. NeuroImage. 2014;100:192–199. 10.1016/j.neuroimage.2014.06.025

23. Rischka L, Gryglewski G, Pfaff S, et al. Reduced task durations in functional PET imaging with [18F]FDG. Neuroimage. 2018;181:323–330. 10.1016/j.neuroimage.2018.06.079

24. Ionescu TM, Amend M, Hafiz R, et al. Elucidating the complementarity of resting-state networks derived from dynamic [18F]FDG and fMRI. Neuroimage. 2021;236:118045. 10.1016/j.neuroimage.2021.118045

25. Deery HA, Liang EX, Siddiqui MN, et al. Reconfiguration of metabolic connectivity in ageing. Commun Biol. 2024;7(1):1600. 10.1038/s42003-024-07223-0

26. Deery HA, Liang EX, Moran C, et al. Metabolic connectivity has greater predictive utility for age and cognition than functional connectivity. Brain Commun. 2025;7(1):fcaf075. 10.1093/braincomms/fcaf075

27. Voigt K, Liang EX, Misic B, et al. Metabolic and functional connectivity provide unique and complementary insights into cognition. Cereb Cortex. 2023;33(4):1476–1488. 10.1093/cercor/bhac150

28. Huang Q, Zhang J, Zhang T, et al. Age-associated reorganization of metabolic brain connectivity in Chinese children. Eur J Nucl Med Mol Imaging. 2020;47(2):235–246. 10.1007/s00259-019-04508-z

29. Vallini G, Silvestri E, Volpi T, et al. Individual-level metabolic connectivity from dynamic [18F]FDG PET reveals glioma-induced impairments. Eur J Nucl Med Mol Imaging. 2025;52(3):836–850. 10.1007/s00259-024-06956-8

30. Sebenius I, Dorfschmidt L, Seidlitz J, et al. Structural MRI of brain similarity networks. Nat Rev Neurosci. 2025;26(1):42–59. 10.1038/s41583-024-00882-2

31. Sebenius I, Seidlitz J, Warrier V, et al. Robust estimation of cortical similarity networks from brain MRI. Nat Neurosci. 2023;26(8):1461–1471. 10.1038/s41593-023-01376-7

32. Fornito A, Arnatkevičiūtė A, Fulcher BD. Bridging the Gap between Connectome and Transcriptome. Trends Cogn Sci. 2019;23(1):34–50. 10.1016/j.tics.2018.10.005

33. Fischl B. FreeSurfer. Neuroimage. 2012;62(2):774–781. 10.1016/j.neuroimage.2012.01.021

34. Glasser MF, Sotiropoulos SN, Wilson JA, et al. The minimal preprocessing pipelines for the HCP. NeuroImage. 2013;80:105–124. 10.1016/j.neuroimage.2013.04.127

35. Jenkinson M, Beckmann CF, Behrens TEJ, et al. FSL. Neuroimage. 2012;62(2):782–790. 10.1016/j.neuroimage.2011.09.015

36. Smith SM, Jenkinson M, Woolrich MW, et al. Advances in functional and structural MR image analysis and implementation as FSL. Neuroimage. 2004;23 Suppl 1:S208–219. 10.1016/j.neuroimage.2004.07.051

37. Hoopes A, Mora JS, Dalca AV, et al. SynthStrip: skull-stripping for any brain image. Neuroimage. 2022;260:119474. 10.1016/j.neuroimage.2022.119474

38. Glasser MF, Coalson TS, Robinson EC, et al. A multi-modal parcellation of human cerebral cortex. Nature. 2016;536(7615):171–178. 10.1038/nature18933

39. Erlandsson K, Buvat I, Pretorius PH, et al. A review of partial volume correction techniques for emission tomography. Phys Med Biol. 2012;57(21):R119–159. 10.1088/0031-9155/57/21/R119

40. Thomas BA, Cuplov V, Bousse A, et al. PETPVC: a toolbox for partial volume correction in PET. Phys Med Biol. 2016;61(22):7975–7993. 10.1088/0031-9155/61/22/7975

41. Alexander-Bloch AF, Shou H, Liu S, et al. On testing for spatial correspondence between maps of brain structure and function. Neuroimage. 2018;178:540–551. 10.1016/j.neuroimage.2018.05.070

42. Mesulam MM. From sensation to cognition. Brain. 1998;121 (Pt 6):1013–1052. 10.1093/brain/121.6.1013

43. Romero-Garcia R, Whitaker KJ, Váša F, et al. Structural covariance networks are coupled to expression of genes enriched in supragranular layers. Neuroimage. 2018;171:256–267. 10.1016/j.neuroimage.2017.12.060

44. Arnatkeviciute A, Fulcher BD, Oldham S, et al. Genetic influences on hub connectivity of the human connectome. Nat Commun. 2021;12(1):4237. 10.1038/s41467-021-24306-2

45. Seidlitz J, Váša F, Shinn M, et al. MSN Detect Microscale Cortical Organization. Neuron. 2018;97(1):231-247.e7. 10.1016/j.neuron.2017.11.039

46. Wang GZ, Belgard TG, Mao D, et al. Correspondence between Resting-State Activity and Brain Gene Expression. Neuron. 2015;88(4):659–666. 10.1016/j.neuron.2015.10.022

47. Burke SN, Barnes CA. Neural plasticity in the ageing brain. Nat Rev Neurosci. 2006;7(1):30–40. 10.1038/nrn1809

48. Goyal MS, Vlassenko AG, Blazey TM, et al. Loss of Brain Aerobic Glycolysis in Normal Human Aging. Cell Metab. 2017;26(2):353-360.e3. 10.1016/j.cmet.2017.07.010

49. Na D, Zhang Z, Meng M, et al. Energy Metabolism and Brain Aging: Strategies to Delay Neuronal Degeneration. Cell Mol Neurobiol. 2025;45:38. 10.1007/s10571-025-01555-z

50. Yang Y, Tang S, Wang X, et al. Eigenmode-based approach reveals a decline in brain structure-function liberality. Commun Biol. 2023;6(1):1128. 10.1038/s42003-023-05497-4

51. Deery HA, Liang E, Moran C, et al. Brain glucodynamic variability is an essential feature of the metabolism-cognition relationship. Proc Natl Acad Sci U S A. 2026;123(3):e2510850123. 10.1073/pnas.2510850123

52. Huang SY, Hsu JL, Lin KJ, et al. Characteristic patterns of inter- and intra-hemispheric metabolic connectivity in MCI and AD. Sci Rep. 2018;8(1):13807. 10.1038/s41598-018-31794-8

53. Kuang L, Zhao D, Xing J, et al. Metabolic Brain Network Analysis of FDG-PET in AD Using Kernel-Based Persistent Features. Molecules. 2019;24(12):2301. 10.3390/molecules24122301

54. Peng L, Zhang Z, Chen X, Gao X. Alternation of the Rich-Club Organization of Individual Brain Metabolic Networks in Parkinson’s Disease. Front Aging Neurosci. 2022;14:964874. 10.3389/fnagi.2022.964874

55. Li W, Tang Y, Peng L, et al. The reconfiguration pattern of individual brain metabolic connectome for Parkinson’s disease identification. MedComm. 2023;4(4):e305. 10.1002/mco2.305

56. Hahn A, Reed MB, Vraka C, et al. High-temporal resolution f-PET/MRI reveals coupling between human metabolic and hemodynamic brain response. Eur J Nucl Med Mol Imaging. 2024;51(5):1310–1322. 10.1007/s00259-023-06542-4

