## Supplementary Materials for "Individual-level metabolic connectivity captures cortical morphology and their coupling strengthens with age"

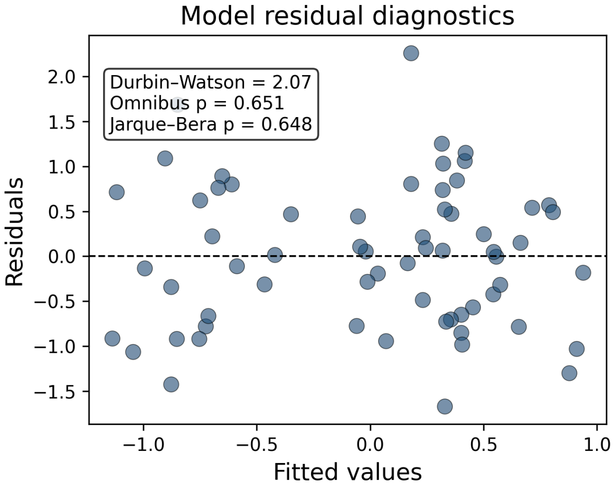


**Fig. S1** Residual diagnostics. Scatter plot of model residuals as a function of fitted values from the robust GLM relating MC–MIND coupling to age, sex, and total intracranial volume. Residuals are symmetrically distributed around zero and show no systematic trends, indicating no major violations of linearity or homoscedasticity. Diagnostic statistics indicate no residual autocorrelation (Durbin–Watson = 2.07) and no significant deviation from normality (Omnibus p = 0.651; Jarque–Bera p = 0.648)

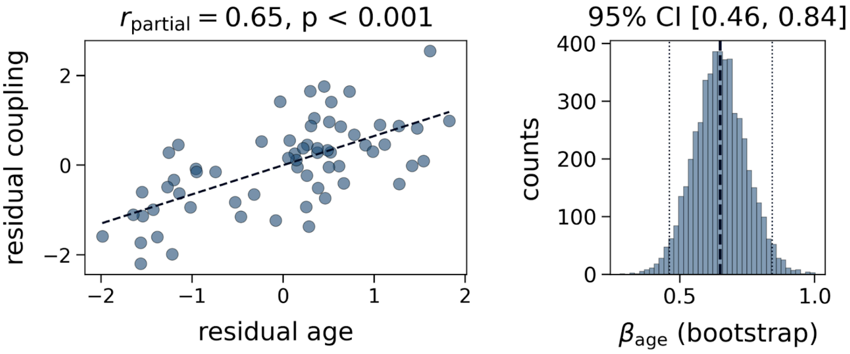


**Fig. S2** Age effect on MC–MIND coupling after partial volume correction. Scatterplot of age versus global MC–MIND coupling, with both variables residualised for sex and TIV. MC was computed using PET data after partial volume correction. The right panel shows the bootstrapped 95% confidence interval (CI) for the regression coefficient. A strictly positive 95% CI indicates a significant age-related increase in MC–MIND coupling independent of partial volume effects

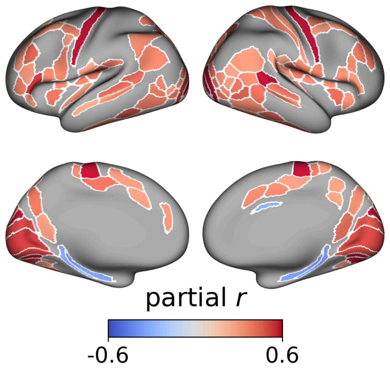


**Fig. S3** Age effects on local MC–MIND coupling after partial volume correction. ROIs showing a significant (FDR-corrected) relationship between local MC–MIND coupling and age, controlling for sex and TIV. MC was computed using PET data after partial volume correction
